# Cortical Surface Area Relates to Distinct Computational Properties in Human Visual Perception

**DOI:** 10.1101/2023.06.16.545373

**Authors:** Scott O. Murray, Tamar Kolodny, Sara Jane Webb

## Abstract

Understanding the relationship between cortical structure and function is essential for elucidating the neural basis of human behavior. However, the impact of cortical structural features on the computational properties of neural circuits remains poorly understood. In this study, we demonstrate that a simple structural feature – cortical surface area (SA) – relates to specific computational properties underlying human visual perception. By combining psychophysical, neuroimaging, and computational modeling approaches, we show that differences in SA in the parietal and frontal cortices are associated with distinct patterns of behavior in a motion perception task. These behavioral differences can be accounted for by specific parameters of a divisive normalization model, suggesting that SA in these regions contributes uniquely to the spatial organization of cortical circuitry. Our findings provide novel evidence linking cortical structure to distinct computational properties and offer a framework for understanding how cortical architecture can impact human behavior.

---

The surface area (SA) of cortical regions varies widely across individuals, even after adjusting for differences in overall brain size. Proportionally greater SA is observed in some regions compared to other regions, and this variation has been associated with behavior in a wide range of cognitive^1-3^ and perceptual^4,5^ domains and clinical conditions^6-12^. However, the impact of these regional SA differences on neural computation and behavior is unknown. We hypothesize that the proportional scaling of SA in a region – determined by whether a region constitutes a relatively large or small portion of the cortex within an individual – affects the organization and spatial properties of neural circuits within that region. We focus our investigation on visual motion processing, examining how individual differences in SA may contribute to variations in perceptual behavior across individuals and the computational mechanisms that may give rise to the specific patterns of behavioral differences.

We measured visual motion duration thresholds – the amount of time a visual motion stimulus needs to be presented in order to perceive its direction. Behavior in this task depends on stimulus size and luminance contrast; paradoxically, larger, high-contrast stimuli require longer presentation durations, reflecting visual surround suppression^13^. We have previously demonstrated that the contrast and size dependency of this perceptual behavior is well accounted for by a divisive normalization model^14^. Divisive normalization describes neural responses in terms of an excitatory drive term, divided by the sum of a spatially broader suppressive drive term, plus a small number, sigma, which controls response sensitivity. Divisive normalization has been used previously to describe responses in a wide range of perceptual and cognitive functions, from early visual processing and attention to olfaction and decision making^15^. Moreover, relevant to our current investigation, we have shown that differences in the spatial width of top-down gain, incorporated into the divisive normalization model, provide a parsimonious account for group differences between autistic and non-autistic participants in this task^16^. Specifically, a narrower width of top-down gain produces a pattern of predicted behavior across size and contrast that approximates the specific pattern of behavioral differences observed in a group of autistic observers versus a non-autistic comparison group. We speculate that individual differences in performance – regardless of autism status – might be at least partially driven by intrinsic differences in the width of top-down gain. In the current study, we explored the potential origin of individual differences in top-down gain as stemming from cortical SA differences in attentional control regions, with a particular emphasis on parietal and frontal regions.

The rationale behind predicting a relationship between SA and the width of top-down gain is based on the spatial organization of attentional control regions (illustrated in Fig. 1). Imagine two individuals with the same total cortical SA, but Person 1 has a proportionally larger attentional control region compared to Person 2. Attentional control regions are known to be spatiotopically organized^17-20^. Assuming that both individuals have identical spatial properties in their feedback circuitry anatomy, we would expect feedback circuits to represent a spatially narrower region of space in Person 1 than in Person 2. This is because an equivalent amount of cortex represents a narrower region of space in Person 1 than in Person 2, leading to a spatially narrower gain amplification in Person 1. To fully explore the implications of this spatial difference in top-down gain, we incorporate it into the normalization model (Fig. 1B) and make behavioral predictions for these two hypothetical individuals (Fig. 1C). Ultimately, we expect that individuals with proportionally larger attentional control regions will demonstrate shortermotion duration thresholds compared to those with smaller attentional control regions, in alignment with these model predictions (Fig. 1C).

**Figure 1.**
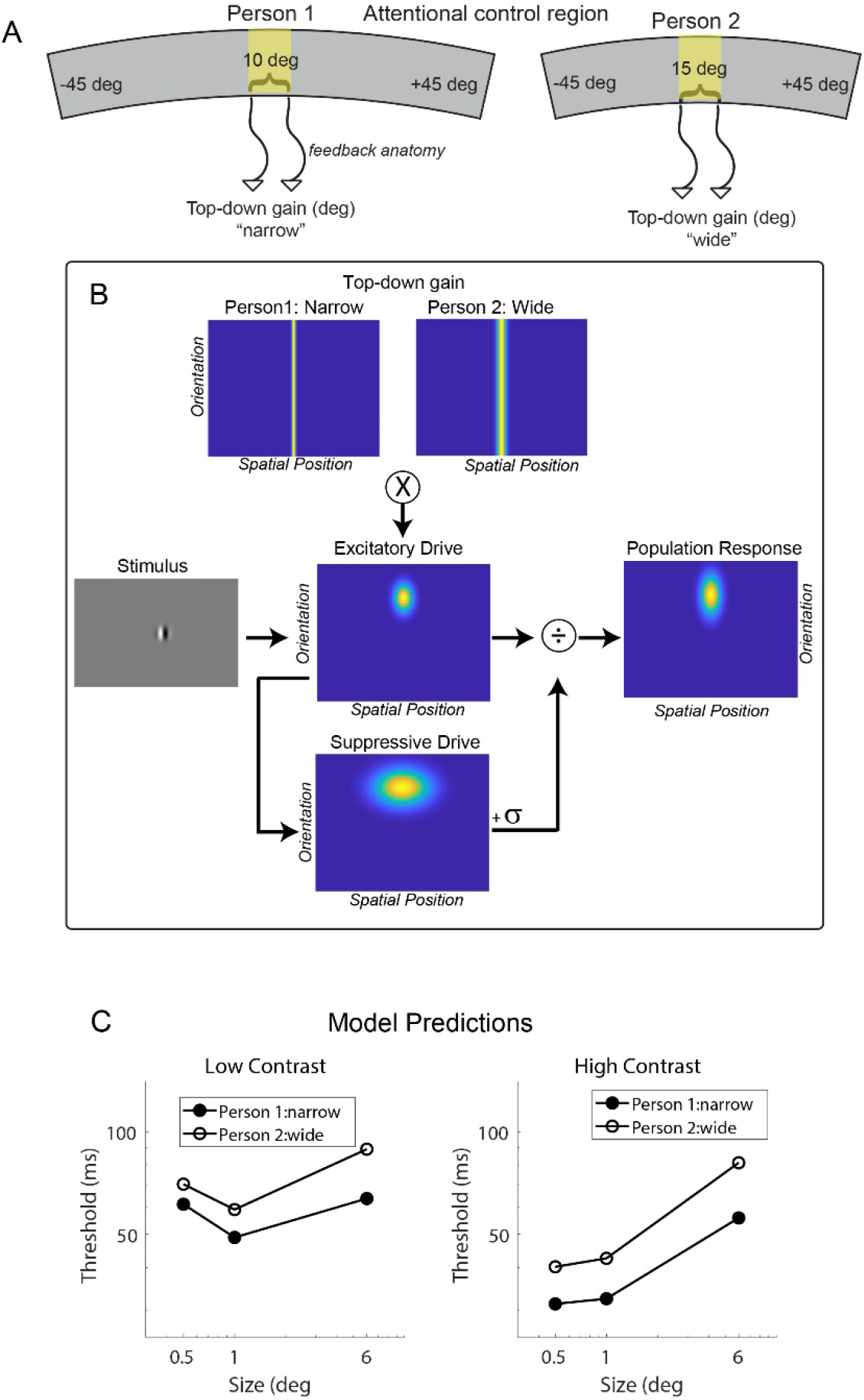
Hypothesized relationship between surface area and its computational and behavioral effects. (A) Schematic illustrating surface area differences between two individuals in a spatiotopically organized attentional control region. An equivalent amount of cortex represents a narrower region of space in Person 1 vs. Person 2. (B) Narrow vs. wide top-down gain incorporated into the normalization model. (C) Predicted duration thresholds from the model for narrow vs. wide top-down gain. Predicted thresholds are derived from the peak population response in the model, assuming higher responses result in shorter thresholds (see Methods).

## Experiment 1: Identifying regions where SA is correlated with duration thresholds

We measured motion duration thresholds using three sizes (stimulus radius = 0.5, 1.0, and 6 degrees) and two luminance contrasts (low = 3% and high = 98%), as in previous publications (Fig. 2;^14,16,21,22^). To test the predicted relationship between SA and threshold, we examined the correlation between individual differences in duration threshold and normalized regional SA, as defined using the Human Connectome Project Multimodal Parcellation atlas (HCP-MMP)^23^, across 62 individuals. The HCP-MMP is comprised of 180 regions per hemisphere that are defined using cortical thickness, myelin content, functional connectivity, and task-based fMRI. To minimize the effects of surround suppression and obtain the most accurate measure of motion duration threshold, we initially focused on behavioral data by averaging thresholds in the two smallest (0.5 and 1.0 deg) high-contrast conditions (later analyses consider all stimulus conditions).

**Figure 2.**
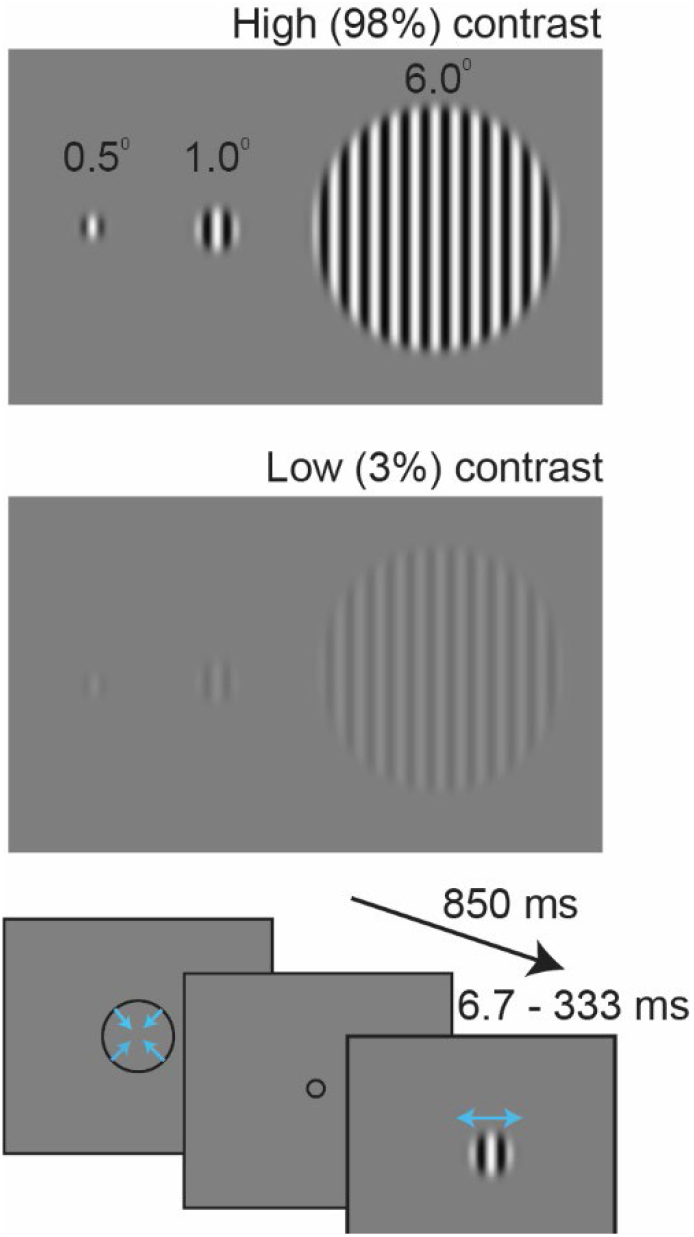
Experimental design. Three different sizes and two luminance contrasts were used. Each trial started with a contracting fixation circle, then a drifting grating was presented and participants indicated whether the motion direction was left or right. Stimuli were presented using an adaptive staircase that adaptively adjusted presentation times to estimate the minimum duration required to perceive leftward or rightward motion (“duration threshold”).

In line with our prediction, significant negative correlations between thresholds and regional SA were observed within well-known attentional control regions (Fig. 3A). Using a statistical threshold of *p* < 0.01 (uncorrected), individual differences in duration thresholds were found to be negatively correlated with individual differences in normalized SA in six parcels of the HCP-MMP atlas – a group of five regions in the right parietal cortex (7AL, VIP, LIPd, IP1, and IP2) and a frontal region in the left hemisphere (11l). Subsequent analyses focus on these “parietal” and “frontal” regions. Relaxing the correlation threshold, for example reducing it to *p* < 0.05, did not substantively alter the regions that correlated with duration thresholds and mostly served to expand the number of identified parietal regions (Fig. 3A, lighter shading). The additional regions in the left anterior temporal pole are discussed in a later separate section.

**Figure 3.**
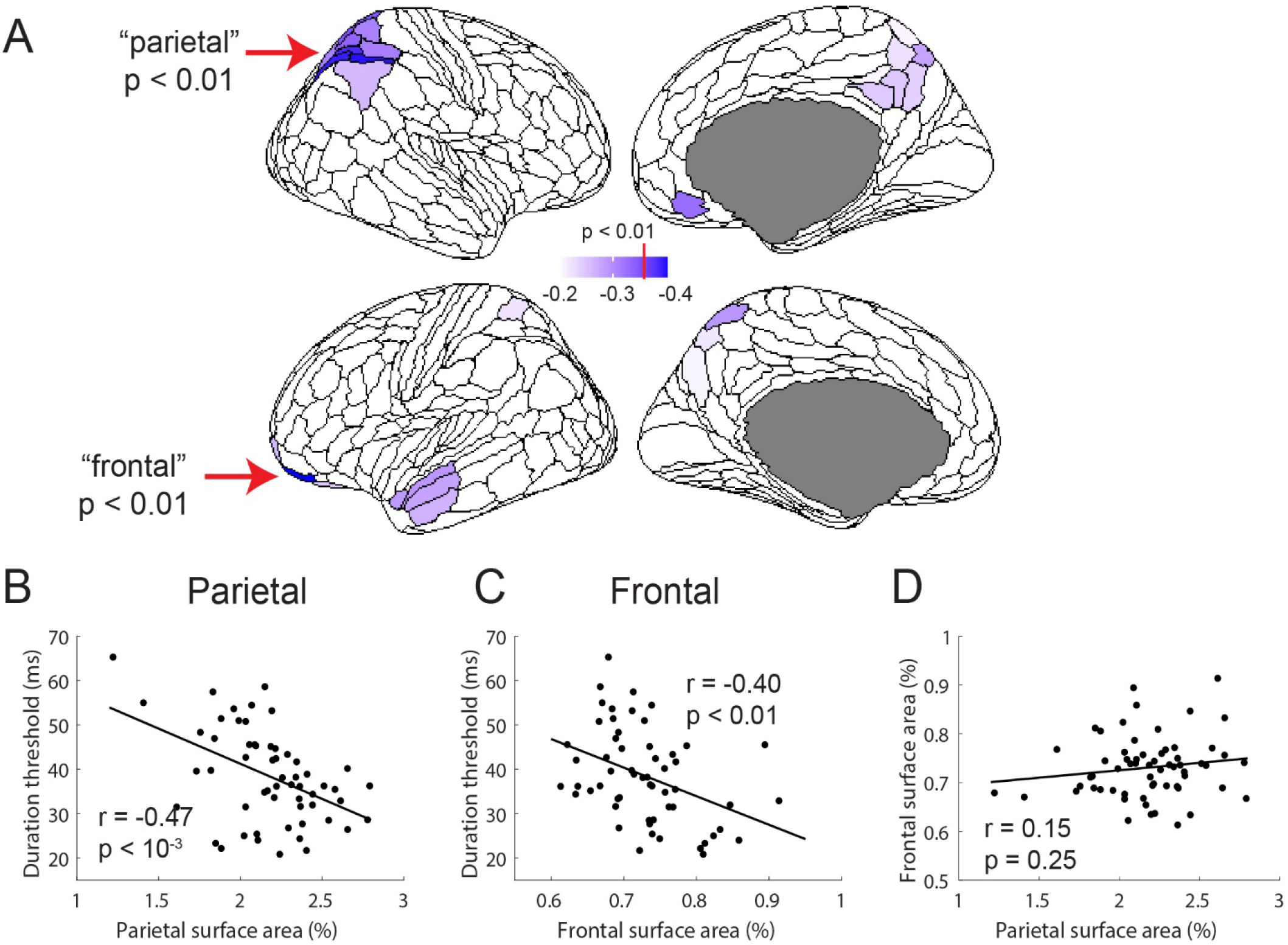
Correlations between surface area and duration thresholds. (A) Regions of the HCP-MMP atlas that negatively correlate with duration threshold. Regions that surpass a p<0.01 are indicated with the red arrows and darker purple color and include 5 parcels in the right parietal cortex and 1 parcel in the left frontal cortex. Significant relationship between individual differences in surface area in the parietal (B) and frontal (C) regions and duration thresholds. (D) There is no significant relationship between individual differences in surface area between the two regions.

While, by definition, the variation in normalized SA in the parietal (summed across the five parcellations) and frontal regions are each significantly negatively correlated with variation in motion duration thresholds (Fig. 3B, *r*_*55*_ = - 0.47, *p* < 10^−3^; and Fig 3C, *r*_*55*_ = -0.40, *p* < 0.01), the variation in SA between the two regions is uncorrelated (Fig. 3D, *r*_*60*_ = 0.15, *p* = 0.25) – a large or small parietal region in an individual is not related to a large or small frontal region. This suggests that SA differences in the parietal and frontal regions may independently contribute to behavior. To explore this possibility, we separately assessed the relationship between normalized SA and behavior for the parietal and frontal regions. First, for a qualitative assessment of the relationship, we divided participants into two groups based on their summed surface area in the parietal parcels. We then plotted the duration thresholds across all stimulus conditions for the two subject groups (small vs. large parietal SA; Fig. 4A). Recall that for the correlational analysis, above, we used the average threshold between the high-contrast 0.5- and 1.0-degree conditions; all of the low-contrast conditions and the large, 6.0-degree, condition are used here for the first time. We performed the same group-split analysis based on normalized SA in the frontal cortex (Fig. 4B).

**Figure 4.**
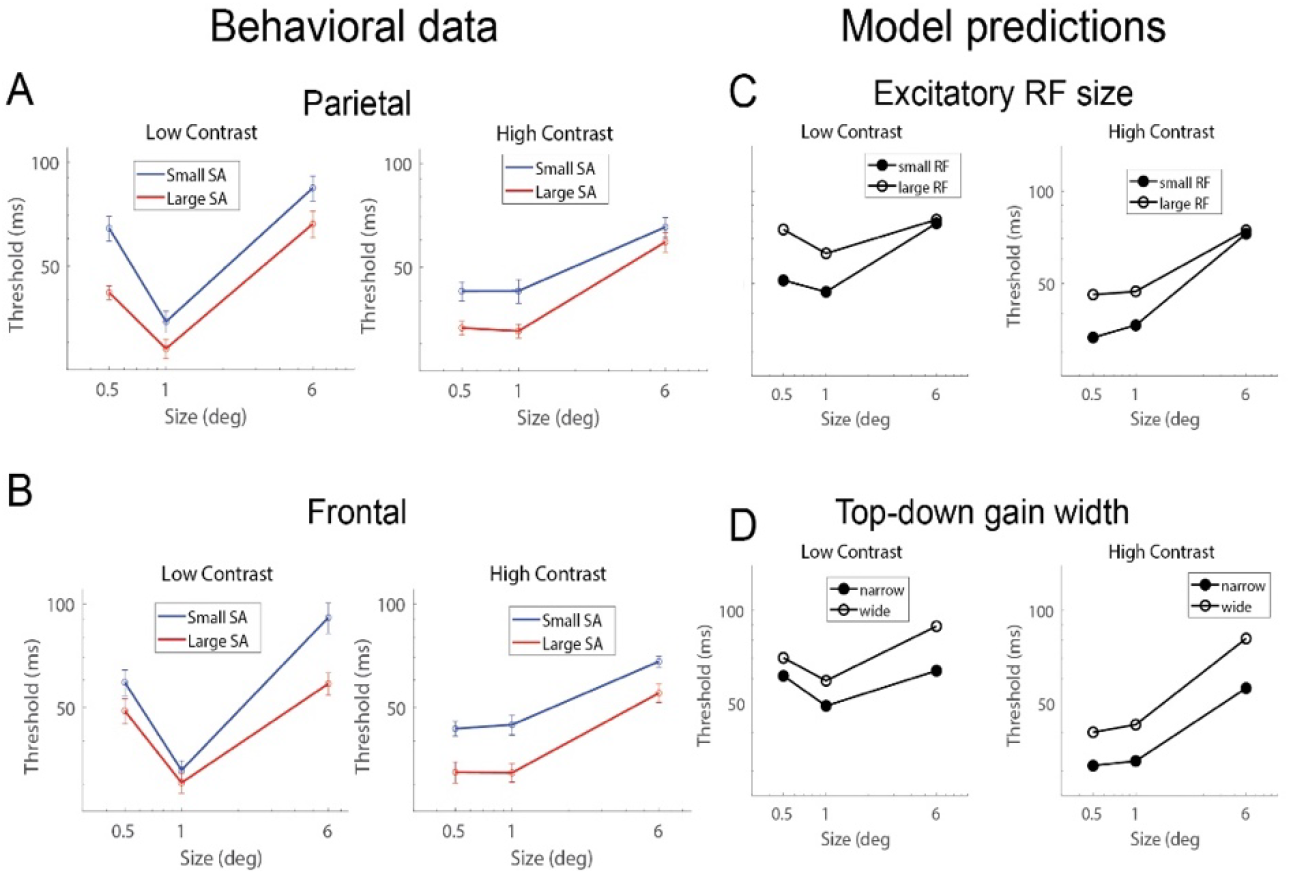
Qualitative analysis. (A) and (B) Subjects were split into two groups – large and small – based on the size of their parietal region and frontal region SA. (C) and (D) Distinct model parameters are associated with the unique behavioral differences across size and contrast. Spatially narrower receptive fields (smaller RFs) produce model predictions that approximate larger parietal SA (A vs. C). Narrower spatial top-down gain produces model predictions that approximate larger frontal SA (B vs. D).

The group-splits revealed a notably differential association with behavior in the parietal and frontal regions, particularly for low contrast stimuli – a marked difference in behavior for the small but not large stimulus when splitting groups based on parietal SA, and a similarly large difference in behavior for the large but not small stimulus when splitting groups based on frontal SA. This differential pattern is further supported by a quantitative analysis – SA in the parietal region correlates with thresholds for the small, low-contrast stimulus but not with the large, low-contrast stimulus (Fig. 5, parietal SA correlated with small, low contrast thresholds, *r*_*55*_ = -0.44, *p* < 0.001; parietal SA correlated with large, low contrast thresholds, *r*_*60*_ = -0.24, *p* = 0.07). Conversely, frontal cortex SA exhibits a significant correlation with the large, low-contrast thresholds but not with the small, low-contrast thresholds (Fig. 5, frontal SA correlated with small, low contrast thresholds, *r*_*55*_ = -0.21, *p* = 0.11; frontal SA correlated with large, low contrast thresholds, *r*_*60*_ = -0.29, *p* = 0.02).

**Figure 5.**
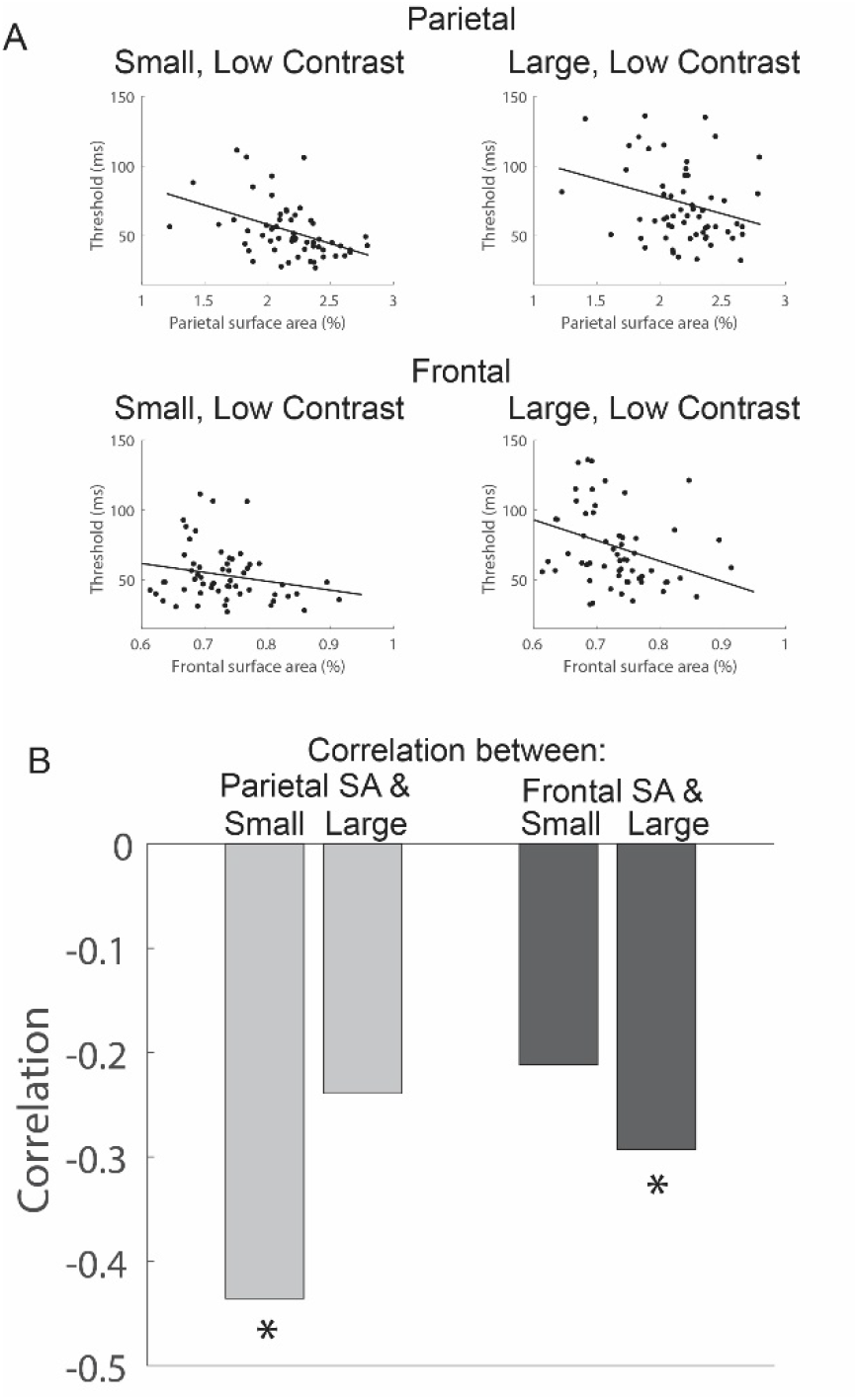
Correlations between motion thresholds for small (0.5 deg) and large (6.0 deg) low-contrast stimuli with SA in the parietal and frontal regions. (A) Individual scatter plots. (B) Correlation values. * p < 0.05

The observed differences in behavior as a function of SA in the parietal and frontal regions can be explained by distinct parameters of the normalization model. Specifically, the pattern of results in the parietal cortex can be approximated by adjusting the spatial extent of receptive fields of the excitatory drive (Fig. 4C) – spatially narrower excitatory drive is predicted to result in lower thresholds for small but not large stimuli. This is because the narrower receptive fields result in less engagement of the suppressive surround. When the stimulus is large, small and large spatial receptive fields engage the suppressive surround equally.

The pattern of behavior linked to frontal cortex is distinct from that associated with the parietal cortex and is more in Fig. 6. In this task, participants performed the motion duration threshold task using a single, small (0.5 deg) high-contrast target stimulus. An uninformative high-contrast surrounding stimulus was also presented, which could move in either the same or opposite direction as the central target stimulus. As a result, participants were encouraged to ignore the surround and focus their attention on the central stimulus. The gap between the surround and target varied from 0.5 to 4.5 degrees.

**Figure 6.**
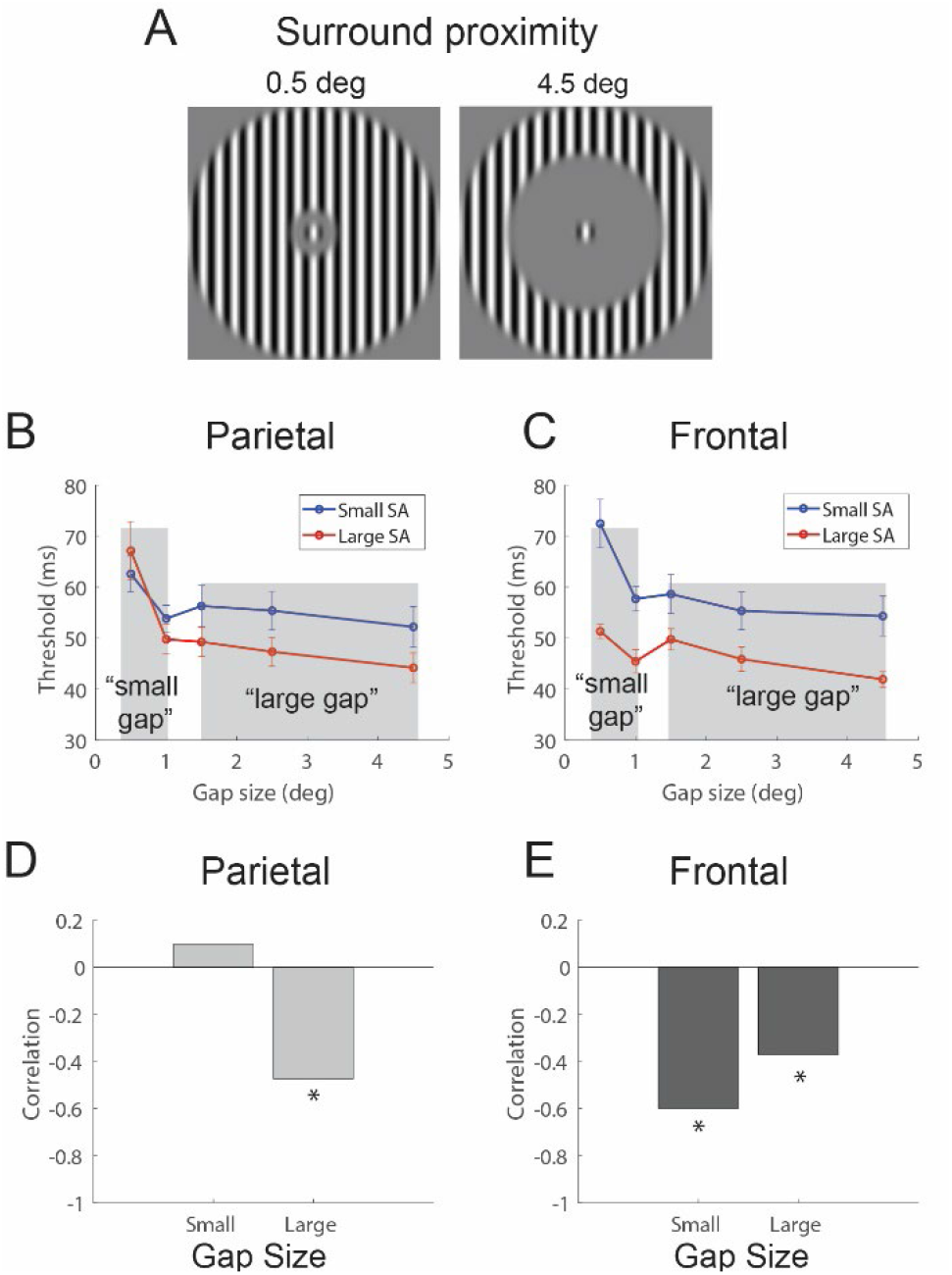
New stimulus configuration. (A) A high contrast small (0.5 deg) central target stimulus was used for the motion duration threshold task. An uninformative high contrast surrounding stimulus could move in either the same or different direction as the central target. The gap size ranged from a minimum of 0.5 deg to 4.5 deg. Subjects were split into two groups based on their SA in parietal (B) and frontal regions. (C) The nature of the group differences across gap size differed. (D) Larger parietal SA was associated with lower thresholds only for large gap sizes. (E) Larger frontal SA was associated with lower threhsolds for all gap sizes.

**Figure 6.**
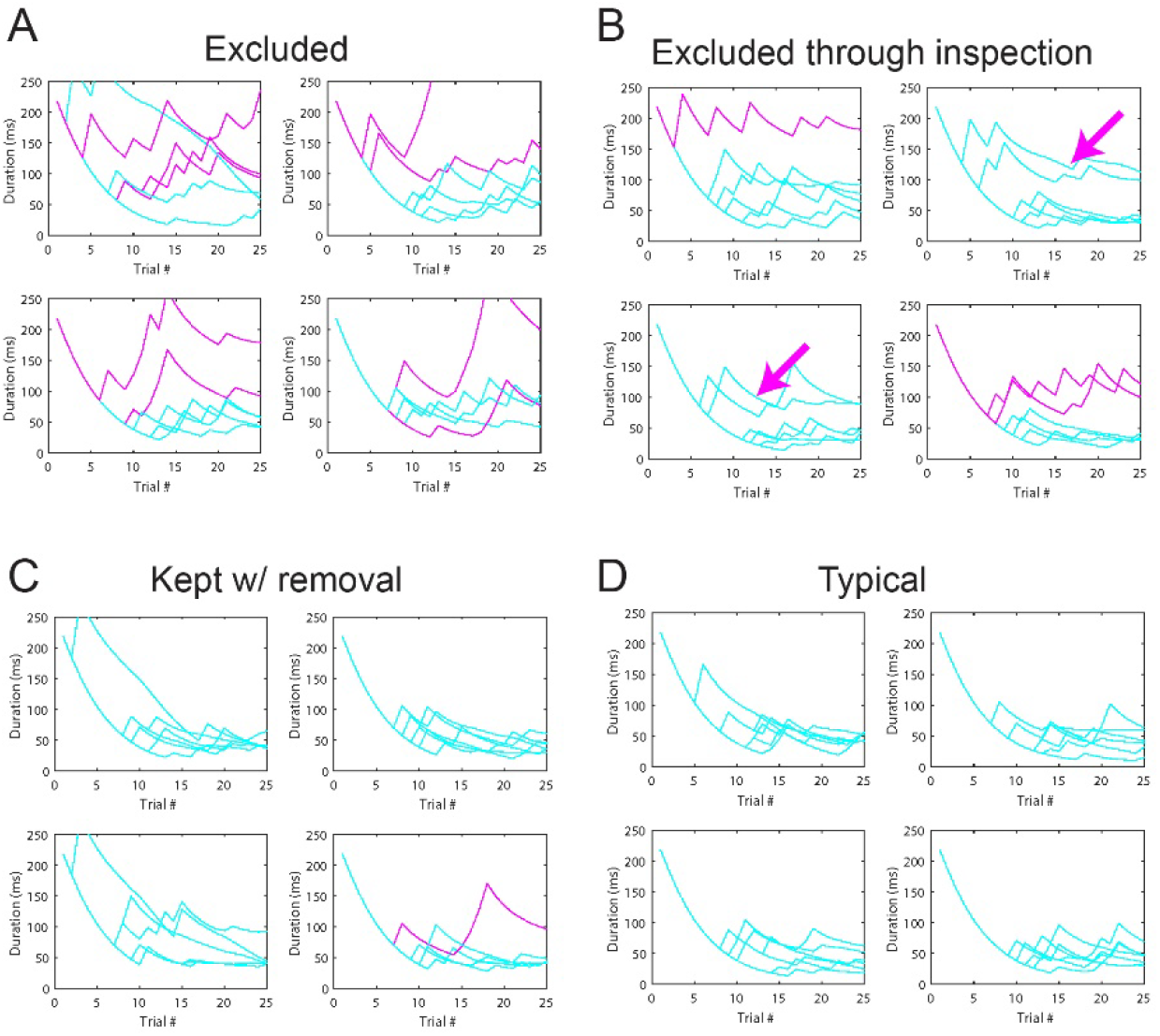
Staircase examples. Each panel is a single run. Each line is a staircase from a particular stimulus condition. (A) and (B) show examples of two of the eleven excluded subjects. Magenta lines are staircases highlighted by the data quality algorithm. (B) Subject was excluded after further visual inspection, noting the 4 staircases with early errors in the trials that prevented convergence along with the 3 magenta, flagged staircases. (C) and (D) are two examples of included subjects. (C) An example of a single staircase (magenta line) that was eliminated from the threshold estimates of an included subject. (D) A typical staircase pattern for a subject with all staircases included.

In the context of the current experiment, this stimulus provides an ideal generalization test for the initial findings. First, consider the predictions for behavioral differences based on SA in the parietal cortex. When the surround is moving in the same direction as the target, as the gap between the surround and target becomes very small, the stimulus essentially resembles a single, large high-contrast stimulus. Conversely, when the gap is large, the stimulus appears as a small, isolated high-contrast stimulus. Predictions for these two extreme conditions can be made by revisiting the results of the parietal cortex group-split for the high-contrast condition in Experiment 1 (Fig. 4A). Specifically, in Experiment 1, small stimuli were associated with parietal cortex SA variation, while large stimuli were not. This same pattern is observed with small and large gap sizes in Experiment 2 – when the gap sizes are very small (approximating a large stimulus), there is no association with SA in the parietal cortex; whereas, when gap sizes are large (approximating a small stimulus), an association is present. This pattern is apparent in both a qualitative analysis based on a group split of closely aligned with our *a priori* hypothesis – it is best accounted for by manipulating the spatial width of top-down gain (Fig. 4D). Spatially wide top-down gain more strongly engages the suppressive surround for large compared to small stimuli. We found no other model parameters that can produce these unique patterns of results for low contrast stimuli. Overall, these findings suggest that SA in these two regions affects normalization-related circuitry in distinct ways.

### Experiment 2: Generalizing the results to a new stimulus configuration

We next aimed to examine the generality of our findings. A subset of the participants took part in a new behavioral experiment, designed to measure individual differences in spatial integration in the motion duration threshold task. The basic configuration of the task is shown parietal cortex SA (Fig. 6B) and a quantitative analysis correlating parietal SA variation with duration thresholds across gap sizes (Fig. 6D). For the correlations, to reduce the number of statistical comparisons and focus on the central question of interest – we averaged thresholds over the two smallest gap sizes (0.5 and 1.0 deg; when the stimulus perceptua **l**y approximates a single, large high contrast stimulus) and the three largest gap sizes (1.5, 2.5, and 4.5 deg; when the central target stimulus is clearly perceptually isolated). Correlating parietal SA with thresholds revealed no relationship for small gap sizes (*r*_*27*_ = 0.10, *p* = 0.62) and a significant relationship with large gap sizes (*r*_*27*_ = -0.47, *p* < 0.01; Fig. 6D).

A different, and much simpler, prediction is made for frontal cortex associations with behavior. Revisiting the results of Experiment 1 (Fig. 4B), larger SA is associated with lower thresholds across all high-contrast stimulus sizes. Similarly, we expect to see an association with frontal cortex SA variation across all stimulus gap sizes in Experiment 2. This prediction holds for both the qualitative group-split (Fig. 6C) and the quantitative analysis of correlations with behavior (Fig. 6E). Specifically, correlating frontal SA with thresholds revealed a significant relationship with small (*r*_*27*_ = -0.60, *p* < 10^−3^) and large (*r*_*27*_ = -0.37, *p* < 0.05) gap sizes. Overall, SA variations in frontal and parietal cortices are associated with distinct patterns of behavior that generalize to new stimulus configurations.

### Experiment 3: Generalizing the results to a new subject cohort

We examined whether the basic findings could be generalized to a new cohort of participants. For that, we reanalyzed an existing dataset of participants diagnosed with ASD who had previously participated in the motion duration threshold experiment and who also had structural MRIs. These participants (n = 20) are a subset of the population we have described in earlier publications; see ^16,24^ for demographic and clinical characterizations. We found that the summed SA in the same six cortical parcellations (parietal and frontal) was associated with duration thresholds in the average of the two smallest (0.5 and 1.0 degree), high-contrast conditions (Fig. 7; *r*_*18*_ = -0.52, *p* = 0.02), demonstrating that the cortical SA model extends beyond our Experiment 1 and 2 discovery sample. Due to the relatively limited number of participants in the Experiment 3 cohort, we did not explore the differential contributions of frontal and parietal cortices.

**Figure 7.**
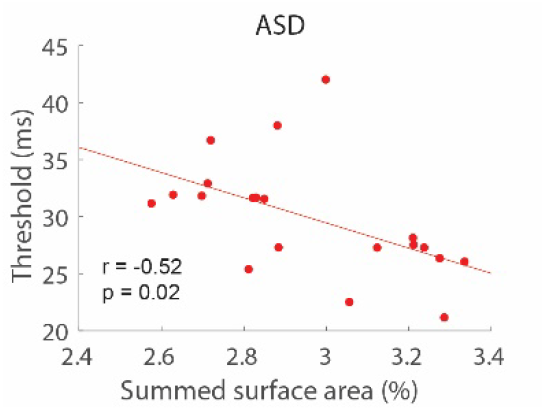
A new cohort. Summed SA across parietal and frontal regions was associated with thresholds in a new cohort of subjects diagnosed with ASD.

### Experiment 4: The relationship between SA and fMRI responses

Our previous research, along with that of others, has demonstrated a strong connection between neural activity in the hMT+ region and performance in motion duration threshold experiments, supporting a key linking hypothesis between neural responses and behavior. Specifically, greater neural response strength in hMT+ is associated with lower duration thresholds. This relationship has been observed using various techniques such as functional MRI (fMRI)^14,25,26^, pharmacology^27^, transcranial magnetic stimulation (TMS)^28^, and monkey electrophysiology^29^. In our earlier studies^22,27^, we found that the degree of hMT+ fMRI response to presentations of drifting gratings correlated with duration thresholds measured in psychophysical experiments performed outside the scanner. This finding indicated that individuals with intrinsically stronger hMT+ neural responses tend to have lower duration thresholds.

Considering this, we investigated whether cortical SA in the identified parietal or frontal regions was related to fMRI responsiveness in hMT+. Our results showed that individual differences in SA within the frontal region (*r*_*28*_ = 0.50, *p* < 0.01), but not the parietal region (*r*_*28*_ = 0.10, *p* = 0.61), were significantly associated with hMT+ fMRI responsiveness (Fig. 8). This further reinforces the idea that the computational architecture in the frontal cortex may contribute to the neural responses underlying task performance.

**Figure 8.**
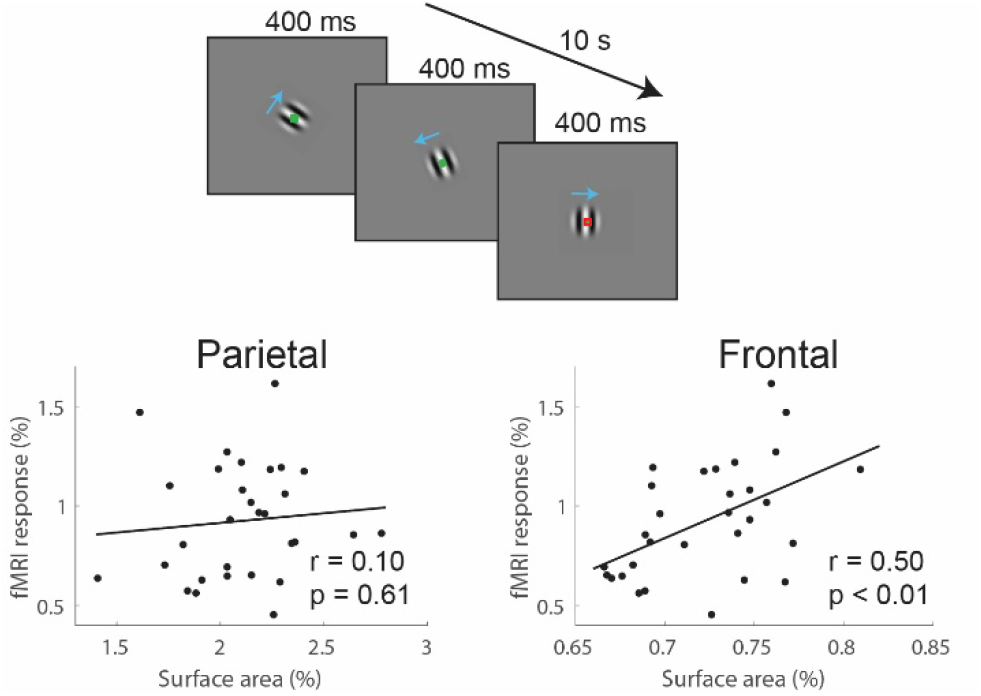
Association with fMRI responses in hMT+. Response magnitude in hMT+ was measured in response to briefly presented small, high contrast gratings. Individual differences in hMT+ responsiveness was associated with SA in frontal but not parietal regions.

### Exploratory analysis 1: No evidence for relationships between visual area SA and behavior

Previous research has demonstrated a link between larger regional SA and improved performance in tasks related to that brain region, suggesting a possible “more cortex is better” hypothesis connecting cortical SA and function. For example, increased SA in V1 is associated with better performance in acuity tasks^4,5^. Consequently, since thresholds are correlated with hMT+ responses^22,27^, we might expect a correlation between SA in motion-sensitive regions like hMT+ and motion duration thresholds. Although our initial analyses did not reveal any significant correlations, it is important to note that extrastriate visual areas, like hMT+, are relatively small and exhibit low positional consistency across individuals, which could impact the strength of observed relationships. This anatomical variability might be responsible for the lack of strong correlations between regional SA in motion-selective regions and motion duration thresholds. Considering these factors, we conducted an exploratory analysis to investigate potential trend-level relationships.

First, we examined the individual correlation values between SA and thresholds in five regions known to exhibit motion-selective neural responses: V1, V3A, V6, MT, and MST. All correlations were positive (i.e., in a direction opposite to the expected relationship), and no p-value was less than 0.14. Next, we explored a different cortical parcellation strategy that focused specifically on visually defined regions in the occipital, temporal, and parietal cortices (“Wang atlas”^30^). We replicated our basic finding using the new parcellation. Specifically, the SA of two right parietal regions of the Wang atlas showed significant negative correlations with duration threshold (right IPS3, *r*_*55*_ = -0.34, *p* = 0.009; right SPL, *r*_*55*_ = -0.34, *p* = 0.01). The SA of known visual motion processing regions such as MT and MST were not associated with duration thresholds (MT, *r*_*55*_ = 0.21, *p* = 0.10; MST, *r*_*55*_ = 0.18, *p* = 0.20). Overall, the results suggest that SA in visual motion processing regions, at least as commonly defined using atlas-based parcellations, are not associated with task performance.

### Exploratory analysis 2: Normalized versus raw surface area

Our hypothesis relating SA to circuitry was originally conceptualized as relating to the proportional enlargement or contraction of SA in a region, rather than the raw surface area of a region. We reasoned that input/output circuitry would simply scale with total brain size, so it was important to account for total surface area in the analyses. Normalizing SA had the added benefit of accounting for differences in total cortical SA that might exist between individuals or groups in our sample (e.g., males vs. females). However, we observed that for many of the relationships we report, normalizing SA may not be necessary. First, normalized (%) and raw (mm) SA are very strongly related (Fig. 9, *r*_*60*_ = 0.76, *p* < 10^−12^). In addition, all of the identified parietal and frontal parcels identified using the correlation between normalized SA and thresholds were also significantly correlated when raw SA area was used. These results did not necessarily need to hold. The fact that normalized and raw SA give similar results suggests that these areas contribute particularly strongly to overall cortical size—that is, if someone has a large cortex, these areas contribute particularly strongly to its size.

**Figure 9.**
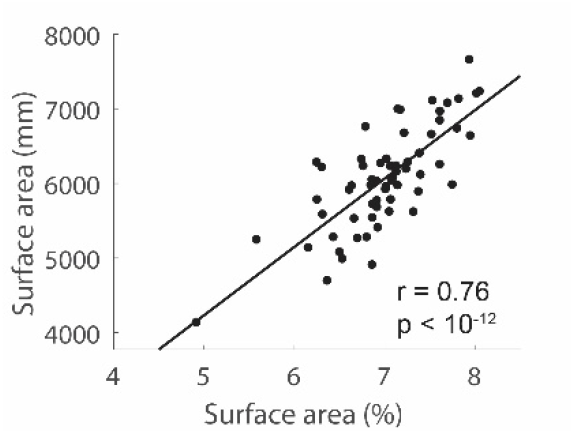
Summed (across parietal and frontal regions) normalized SA (%) and raw SA (mm’s) are highly correlated.

### Exploratory analysis 3: Left anterior temporal pole

For completeness, we report an observed relationship between SA and duration thresholds in a region where we did not have strong *a priori* hypotheses: the left anterior temporal pole. This region is visible in Fig. 3A and includes parcels STGa, STSda, TE1a, and STSva. The relationship for any given parcel was modest, with *r*-values ranging from -0.27 to -0.29 and *p*-values from 0.03 to 0.05. The relationship between SA variation in the temporal pole did not extend to the alternate stimulus configuration (Experiment 2), the additional cohort of participants (Experiment 3), or the fMRI responses (Experiment 4). Therefore, we do not discuss its potential involvement in more detail other than to note it as a potential region of interest for future studies.

## Discussion

Previous studies have shown substantial individual variation in SA in cortical subregions^4,5,31^ and we hypothesized that this variation may be related to the spatial properties of cortical circuits. We tested this hypothesis by measuring the relationship between surface area (SA) in cortical subregions and motion duration thresholds. Our results demonstrate that individuals with proportionally larger subregions of parietal and frontal cortices exhibit shorter motion duration thresholds compared to those with smaller regions. Our results suggest unique computational implications of SA in parietal and frontal cortices, with each region associated with different aspects of the normalization model. A cluster of subregions in the right parietal cortex were associated with the spatial extent of excitatory receptive fields; incorporating spatially narrower excitatory receptive fields into the normalization model produced a predicted pattern of thresholds across size and contrast that was similar to that exhibited by individuals with larger SA in these regions. In contrast, a subregion of the left frontal lobe was associated with the width of top-down gain; incorporating spatially narrower top-down gain into the normalization model produced a pattern of predicted thresholds across size and contrast that was similar to individuals with larger SA in this region.

Although we expected *a priori* to find a relationship between parietal cortex SA and thresholds, we did not expect it to be associated with the spatial extent of receptive field size. In fact, our original conceptualization was that the parietal cortex would be associated with the width of top-down gain, as depicted in Fig. 1. The spatial width of excitatory receptive fields would seem to be a property more appropriate for regions directly involved in sensory processing, such as hMT+ or possibly V1. One possibility is that SA is a feature that represents ‘inherited’ properties of its inputs— that the effects of SA on neural computation is more aligned with input than output properties of the region. This possibility is diagramed in Fig. 10 which shows how the excitatory receptive fields of inputs to parietal cortex could be reflected in differences in spatial activation – narrow versus wide for large versus small SA, respectively. Future work will be required to clarify this level of computational specificity and the degree to which SA is more aligned with spatial properties of inputs, outputs, or some combination of the two.

**Figure 10.**
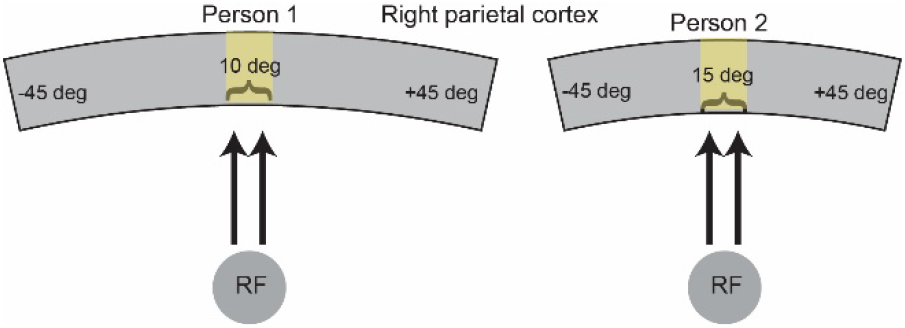
An alternative to the original schematic that highlights the relationship between SA and inputs to a region. Equivalent excitatory inputs would be expected to result in narrow versus wide excitatory drive for large (left) versus small (right) SA.

We did not have strong *a priori* assumptions about the specific contribution of frontal regions. While frontal regions are known to be critical in top-down attentional modulation, the degree to which they are spatiotopically organized is unclear. Thus, in our original hypothesis—depicted in Fig. 1—it was unclear whether a region anatomically so far removed from sensory processing would be associated with visual motion processing. However, as the results clearly support—both in the significant relationship with behavior and with the association with fMRI responsiveness in hMT+— subregions of the frontal cortex appear to be strongly associated with visual sensory processing and in a manner consistent with top-down gain within the normalization model. Again, the relationship between SA and circuitry may be more relevant to the input than outputs of the frontal region.

While the relationship between visual processing and parietal cortical regions is well-established, featuring clear functional connections to visual processing areas in the occipital-temporal cortex^32-34^, the functional connections of the frontal region (left 11l) we identified are less well-known. However, recent connectomic analyses of the HCP data using the HCP-MMP atlas suggest that region 11l is particularly well-suited to modulate visual responses, either indirectly through strong functional connectivity to IP2 in the parietal lobe (one of the regions where SA correlated with thresholds) or more directly through anatomic connections to V1 via the inferior fronto-occipital fasciculus^35^. In fact, recent findings have suggested that the inferior fronto-occipital fasciculus may play a key a role in tuning early visual processing for efficient task performance^36^.

An additional indication that the association between SA variation and motion duration thresholds is related to spatial attention can be observed in the strong right hemisphere bias of the parietal cortex associations. The right parietal cortex’s involvement in spatial attention is well-established across a broad range of methodologies. For example, lesion studies have demonstrated that patients with damage to the right parietal cortex often exhibit spatial neglect, a condition characterized by an inability to attend to or be aware of stimuli in the left side of their visual space^37,38^. This phenomenon is more pronounced and persistent when the lesion occurs in the right hemisphere compared to the left hemisphere^39^. Functional fMRI studies have consistently shown that the right parietal cortex is activated during tasks requiring spatial attention such as orienting, target detection, covert attention shifts, and spatial working memory^40-43^. This asymmetry is further supported by findings that the right parietal cortex exhibits greater functional connectivity with other brain regions involved in attention and perception, such as the frontal eye fields and extrastriate visual cortex^44,45^. Thus, while the motion duration threshold task does not directly manipulate the focus of spatial attention, the fact that individual differences in performance can be well-explained by manipulating the top-down gain parameter of the normalization model and that the relationship between SA variation in parietal cortex and thresholds are strongly right lateralized is consistent with the hypothesis that these associations reflect underlying neural processes related to spatial attention and its modulation by cortical structure.

Interestingly, our findings were not consistent with a simple “more cortex is better” hypothesis. Previous research has demonstrated a positive association between SA in regions typically thought to play a role in a behavior and performance in that behavior. For example, individuals with larger SA in V1 exhibit lower thresholds in acuity tasks^4,5,46^. Although it is evident that responses in hMT+ are crucial for performance in this task^14,28,29^, we did not observe a relationship between thresholds and SA in visual motion processing regions, such as MT and MST, at least when defined using common atlas-based parcellations. Therefore, while activity in hMT+ may be essential for task performance, it seems that modulation of activity in hMT+ through top-down influences may be critical for determining individual differences in performance.

Our results demonstrate the predictive ability of our findings, as we observed similar associations between cortical SA and duration thresholds in a second task using a new stimulus configuration, as well as in an additional cohort diagnosed with autism spectrum disorder (ASD). This indicates that the relationship between cortical SA and motion duration thresholds extends beyond our initial sample, highlighting the potential of using structural MRI measures to predict task performance in various populations. For example, previous research has demonstrated a variety of (and sometimes conflicting) findings with autistic participants in this task^16,21,47,48^. This has led to differing computational accounts and speculations about how neural processing may be altered in ASD. The current findings point to possible neuroanatomical bases for these conflicting results – SA differences appear to contribute to performance differences in computationally unique ways, depending on the region. Consequently, when considering autistic participants, it may be necessary to characterize subpopulations with differences in SA in parietal cortex and other subpopulations with differences in frontal cortex. In fact, some previous accounts align precisely with the computational roles attributed to parietal and frontal regions – one model of differences found in ASD emphasized a change in excitatory receptive field width^48^, while another account suggested changes in the width of top-down gain^16^. Overall, accounting for the specific variations in SA may be crucial for understanding how the task may differ in those with autism and other clinically defined groups. Unfortunately, our current limited sample size of autistic participants restricted our ability to explore these issues. Future studies that incorporate larger sample sizes that include both structural and psychophysical measures will help clarify this issue.

One potential limitation of our study is the reliance on atlas-based parcellations to define regions of interest. These predefined regions may not fully capture individual variability in the location and size of attentional control and motion processing regions. One possible solution to address this limitation is the use of individualized, targeted functional localizer tasks for defining regions. However, implementing this approach in a large sample would require considerable resources to specify multiple ROIs. An alternative strategy could involve exploring multiple, automatic parcellations to first determine the appropriate spatial scale of SA variation. Although the optimal spatial scale remains unclear, our results were consistent across the regions defined by the HCP-MMP atlas and the visual cortex-optimized Wang (2015) atlas^30^. Future research should focus on examining SA variation across multiple spatial scales to establish the optimal spatial scale for characterizing SA in the context of brain-behavior associations.

## Methods

### Participants

The inclusion criteria for participants in all experiments were as follows: age 18-30 years, non-verbal IQ >70, normal or corrected to normal visual acuity, no visual impairments, no impairment to sensory or motor functioning, no history of seizures or diagnosis of epilepsy, and no neurological disease or history of serious head injury. A total of 73 participants participated in Experiment 1, with a subset of 30 of these participants described in previous publications^14,22^. Due to poor psychophysical data quality, 11 participants were excluded from the analysis as their staircases failed to estimate a reliable threshold. These participants were identified and removed through a combination of an automatic algorithm (described in the psychophysics section below) and visual inspection. Consequently, the results of a total of 62 participants are reported in Experiment 1. For five participants, thresholds for the smallest (0.5 deg) condition were not measured, so when reporting statistical analyses that include this condition, 57 participants are included. All other analyses include the full set of 62 participants.

Experiment 2 (alternative stimulus configuration) included 32 participants from the original 73 participants who participated in Experiment 1. The psychophysical data from three of these participants were excluded from analysis due to poor psychophysical data quality.

|  | N | M/F/NS* | Age (mean, SD) |
| --- | --- | --- | --- |
| **Experiment 1:**<br>**Psychophysical motion**<br>**duration threshold task** | 62 | 35/27 | 23.4 (3.4) |
| **Experiment 2:**<br>**Psychophysical spatial**<br>**integration task** | 29 | 14/13/2 | 23.1 (3.0) |
| **Experiment 3:**<br>**ASD cohort** | 20 | 12/8 | 22.5 (3.7) |
| **Experiment 4:**<br>**fMRI response in hMT+** | 30 | 18/12 | 23.3 (3.5) |
| Note: The cohorts of experiments 2 and 4 are non-overlapping sub-samples of the cohort of experiment 1. The number of participants reported in the table is after data-quality exclusions. NS = not specified |  |  |  |

The participants in experiment 3 were individuals with ASD, who were included in previous publications^16,24,49^. Full diagnostic characterization can be found in these previous reports. A total of 23 autistic individuals had complete data across the necessary stimulus conditions, and three were removed using the same data quality exclusion procedures, resulting in a total of 20 subjects.

A portion of the fMRI data in experiment 4 have been described previously^22,27,49^ and represent a subset of the subjects that also had psychophysical data (n = 30).

### MRI acquisition

MRI data of experiments 3 & 4 were acquired using a Philips 3-Tesla Achieva scanner. MRI data of the remaining 43 participants in experiment 1 (including the participants of experiment 2) were acquired using a Philips 3-Tesla Ingenia Elition scanner. In Both scanners, a 32-channel high-resolution head coil was used. High resolution T1-weighted anatomical images were acquired for each participant using a magnetization prepared rapid acquisition gradient echo (MPRAGE) protocol (TR = ∼7.7 msec, TE = ∼3.5 msec, flip angle = 7°, 176 1 mm thick slices, 256 × 256 × 176 matrix, voxel resolution: 1 × 1 × 1 mm). There were no differences in SA between participants scanned in the different scanners.

### Cortical Surface Area

Cortical surface area was measured using FreeSurfer(7.2.0). The HCP-MMP atlas was projected onto the cortical surface of each subject using FreeSurfer’s mri_surf2surf command and the lh/rh.HCP-MMP1.annot annotation file. Statistics, including surface area (mm), were generated using FreeSurfer’s mris_anatomical_stats command. Unless noted, all surface area values were normalized to the total cortical surface area per hemisphere (for example, RH_V1_normalized = (RH_V1 / (sum(all_RH_parcels))) * 100). With 180 parcels per hemisphere, each region comprised roughly 0.5 to 1.5% of the total hemispheric cortical surface area. To extract surface area for the Wang et al. (2015) atlas, the Python library <u>Neuropythy</u> was used with its “atlas” function. Once applied, the Python <u>Nibabel</u> library was used to extract surface area values. These values were normalized using the total cortical surface per hemisphere derived from the FreeSurfer atlas. Visualization (Fig. 3) was done using the R package ggseg^50^.

Grouping subjects based on large and small total cortical SA was achieved by first calculating the sum of normalized SA in the parietal/frontal cortex regions that correlated with duration thresholds. Next, the top 1/3 and bottom 1/3 of subjects in total SA were selected to form the groups (Fig. 4A and 4B). For experiment 2 (Fig. 6B and 6C), groups were created using the top and bottom halves of subjects, due to the smaller overall number of participants.

### Psychophysics. Visual display and stimuli

Our experimental apparatuses and stimuli have been described in our recent publications^14,16^. We used a ViewSonic PF790 CRT monitor (120 Hz) and Bits# stimulus processor (Cambridge Research Systems, Kent, UK). Stimuli were created and displayed in MATLAB (MathWorks, Natick, MA) and PsychToolbox^51-53^. Viewing distance for all experiments was 66 cm, and luminance was linearized using a PR650 spectrophotometer (Photo Research, Chatsworth, CA). Stimuli were sinusoidally modulated luminance gratings presented on a mean luminance background. Vertically oriented gratings drifted either left or right (drift rate = 4 cycles/ s) within a circular aperture, which was blurred with a Gaussian envelope (SD = 0.21°). We used three different stimulus sizes: 0.5°, 1.5°, and 6° in diameter. The Michelson contrast of the gratings was either 3% (low) or 98% (high), and the spatial frequency was 1.2 cycles/°.

### Psychophysics. Experiment 1 & 3, Motion duration thresholds

Participants discriminated the direction of motion (left or right) of a briefly presented drifting grating. Trials began with a shrinking circle fixation mark (850 ms) at the center of the screen, followed by a vertical grating, and then a response period (no time limit). Grating duration was adjusted across trials (range 6.7–333 ms) according to an adaptive (Psi) staircase procedure implemented with in the Palamedes toolbox^54^. Correct responses tended to yield shorter durations on subsequent trials, adjusting the grating duration to find the briefest presentation for which the participant would perform with 80% accuracy. Each staircase consisted of 25 trials. Six independent staircases (3 sizes × 2 contrasts) were included in each run and were randomly interleaved across trials. A total of four runs were included in each experimental session, which began with examples and practice runs to familiarize subjects with the procedure. Total task duration was approximately 30 minutes.

Duration thresholds for motion direction discrimination were calculated by fitting a Weibull function to the data from each individual staircase. Guess rate and lapse rate were fixed at 50% and 4%, respectively. Thresholds were calculated from the fit psychometric function as the duration value where the participant performed with 80% accuracy. The median of the four threshold estimates was used as the final threshold estimate for a particular condition.

Because individual differences in thresholds are central to the analyses, extra care was taken to ensure accurate threshold estimates for each subject. Staircase quality was determined by calculating the ratio of the threshold estimates across trials. The average estimate of the final 5 trials of the staircase was divided by the average estimate of the first 10 trials. If this ratio exceeded 0.8, the staircase was removed from the average. This procedure detects “rising staircases” near the end of the trial sequence and was confirmed via visual inspection to identify unreliable estimates. A subject with 4 or more flagged staircases was considered for exclusion. For excluded subjects, the algorithm identified, on average, 5 staircases per subject for removal. For included subjects, the algorithm identified, on average, 1 staircase per subject. However, the algorithm sometimes failed to detect staircases when errors occurred near the beginning of the trial sequence; these errors result in particularly high threshold estimates at the beginning of the sequence. These cases were determined through visual inspection. Figure 11 shows examples of different situations for four subjects: A) an excluded subject, B) a subject excluded after visual inspection (a case where the algorithm failed to detect several poor staircases), C) a subject included but with a single staircase removed from the final estimate, and D) a typical subject with all data included.

### Psychophysics. Experiment 2, Motion integration

Experiment 2 was the same as Experiment 1 in all respects except that it used a single, small (0.5 deg) target grating and a surrounding grating that could move in either the same or opposite direction as the target. The gap size between the outer edge of the target and the inner edge of the surround varied, including 0.5, 1.0, 1.5, 2.5, and 4.5 degrees. The widest portion of the surrounding grating extended to 7.75 degrees. Trials were also included that had no surround. One staircase per stimulus condition was included in each of four runs, and staircases were interleaved across trials. Each staircase was composed of 25 trials. In this report, only data from the 5 gap sizes for the same-direction surround are presented.

### Functional MRI. Experiment 4, hMT+ responsiveness

Portions of the fMRI data have been described previously^22,27,49^ and are a subset of the sample in Experiment 1. Briefly, the fMRI measurements were designed to assess individual differences in responsiveness in hMT+ to stimuli that approximated the stimuli used in the psychophysical experiment. As such, a block-design was used that varied stimulus contrast across blocks (3%) and (98%). The scans began with a blank block in which only the fixation mark was presented on a mean gray background. Blocks of drifting gratings (2° diameter) were then presented centrally in an alternating order, each followed by a blank block, to permit the fMRI response to return to baseline (6 low-contrast blocks, 6 high contrast, and 13 blank per run). During a stimulus block, drifting gratings of different direction were presented for 400 ms. Subjects each completed 2–4 runs of the contrast experiment. Throughout all blocks in all functional scans, participants were engaged in a conjunctive color-shape detection task in the center of the screen, to encourage fixation and maintain engagement and wakefulness. Participants were instructed to press a button when a green circle appeared in a series of small, briefly presented color shapes.

Functional localizer scans were acquired to identify the hMT+ regions-of-interest (ROI). To identify human MT complex (hMT+) stimuli were sinusoidal luminance modulated gratings at 15% contrast. Stimuli were the same size and had the same spatial frequency as the experimental stimuli, and were presented in the same retinotopic locations, without edge blurring. Blocks of drifting gratings (4 cycle/°, same drift rate of the experimental stimuli) alternated with blocks of static gratings, each block 10 s long, for a total of 24 blocks per scan.

Data were preprocessed using BrainVoyager QX v2.8.4 (Brain Innovation) software. EPI data were motion-corrected, corrected for distortion due to magnetic field inhomogeneities, high-pass filtered, and coregistered to the AC– PC aligned T1 structural scan. ROIs were identified from the localizer scan data using correlational analyses with an initial threshold of *p* < 0.05 (Bonferroni corrected). ROIs were defined for each hemisphere in the anatomical region of motion-selective MT in the lateral occipital lobe. Average time courses across the 20 most significantly active (from the localizer scan) voxels in each ROI were determined for each block. Percentage-transformed time courses were then calculated for each block: first, for each stimulus block, we extracted 12 time points corresponding to -4 s before stimulus onset to 18 s after stimulus onset. Then we converted the values to percentage signal change relative to the mean value of the three time points before stimulus onset. To extract a univariate measure to be used in the statistical analyses, the average response magnitude in time points 8–12 s was calculated from each of these time courses. Some blocks/runs were excluded from averaging because of head motion or low task performance, according to the following criteria: data for a given block was excluded because of head motion if the framewise displacement in successive TRs was >0.9 mm^55,56^, up to and including 8 TRs before the stimulus block or 2 TRs after it. If more than one-half the blocks of either condition were excluded, or a subject had a hit rate of <60% in the central fixation task, the whole run was excluded.

### Computational Model

The normalization model used in this study is the same as in recent publications^14,16^ and is based on the instantiation first described by Reynolds and Heeger^57^. The model is useful for making qualitative comparisons between versions using different parameters. It is employed to make predictions in the motion duration threshold task by assuming that thresholds are inversely proportional to response magnitude – a higher neural response in the model results in a lower predicted duration threshold. The scalar value used as the model response is the maximum value of the population response image. A scaling factor, applied across all stimulus conditions, is used to convert the model response to milliseconds. The comparison of differences in the width of top-down gain used the same model as reported previously to account for differences between autistic and non-autistic subjects in this same motion duration threshold task^16^. The version used to characterize differences in the spatial width of excitatory drive held all other parameters constant and used values of 3 and 4.5 for the excitatory drive widths (arbitrary units). The version used to characterize differences in the attention width used values of 3 and 7 (arbitrary units). MATLAB code is available upon request that implements these models. Since the primary hypothesis is that cortical surface area affects spatial circuitry, we also explored the only other spatial characteristic of the model: the spatial pooling width of the divisive normalization. This did not yield predictions that were related to the observed pattern of behavior as a function of SA differences, only serving to additively shift predicted thresholds.

## Acknowledgments

This research was supported by NIH R01MH11847 to S.O.M. and S.J.W. and R01 MH106520 to S.O.M. We thank Noah Benson for assistance with the implementation of Neuropythy.

